# Fertilizer reduction with bio-organic fertilizer to regulate the root soil microbial community structure to improve Baby Chinese cabbage yield

**DOI:** 10.1101/2022.02.28.482435

**Authors:** Li Jin, Ning Jin, Shuya Wang, Jinwu Li, Xin Meng, Yandong Xie, Yue Wu, Shilei Luo, Jian Lyu, Jihua Yu

## Abstract

Using high-throughput sequencing, this study aimed to explore the response of soil microbial community and Baby Chinese cabbage yield to the reduction of chemical fertilizers combined with bio-organic fertilizer in the Gansu plateau, China. Our experiments consisted of conventional fertilizer (CK), 30% chemical fertilizer reduction + 6,000 kg bio-organic fertilizer (T1), 30% chemical fertilizer reduction + 9,000 kg bio-organic fertilizer (T2), 40% chemical fertilizer reduction + 6,000 kg bio-organic fertilizer (T3), and 40% chemical fertilizer reduction + 9,000 kg bio-organic fertilizer (T4). Compared with CK, soil microbial diversity and richness were higher for all treatments with added bio-organic fertilizer. PCoA showed that the bacterial and fungal communities in T2 and T4 were similar to each other. Redundancy and Spearman’s correlation analyses of microbial communities and soil physicochemical properties revealed that reductions in chemical fertilizer rate combined with bio-organic fertilizer had a stronger impact on the fungal than the bacterial community. They also increased the relative abundance of the dominant bacterial and fungal phyla. Baby Chinese cabbage yield was relatively higher under the combined bio-organic fertilizer plus reduced chemical fertilizer rate with T2 showing the highest yield. Therefore, this approach is feasible for sustainable agricultural, cost-effective and profitable crop production.

**Importance:**

- Bio-organic + moderately reduced chemical fertilizer raised Chinese cabbage yield
- Bio-organic + chemical fertilizer was more efficacious than either one alone
- Presence of bio-organic fertilizer enhanced overall rhizosphere physicochemistry
- Bio-organic fertilizer improved beneficial bacterial & fungal abundance & diversity
- Fertilizer combination sustainably & cost-effectively improves crop & soil quality

## 1. Introduction

Gansu Province, China is at high altitude, has abundant light, experiences wide diurnal temperature differences, and has low summertime temperatures, which makes cold-grown vegetable crops that are rich in nutrients and have excellent flavor. This area is dry and has low precipitation rates and few diseases and insect pests. Pesticide use is minimal in this region; thus, crop quality and safety have been assured. Gansu Plateau summer vegetables have bright colors, appealing shapes and flavors, and high national consumer demand. Baby Chinese cabbage (*Brassica rapa subsp. Pekinensis*) is a subspecies of Brassica oleracea in the crucifer family. This crop prefers cold climates and has become a mainstay of the summer vegetables in Gansu Plateau. Its planting scale is expanding annually. To contend with growing crop demand and improve yield, traditional agricultural systems have relied upon high application rates of chemical fertilizer (Hartmann et al. 2015). Chemical fertilizer overuse is also accompanied by a reduction in soil nutrient absorption efficiency, soil quality deterioration, greenhouse gas emissions, and aquatic ecosystem eutrophication (Zhu et al. 2016, Ajeng et al. 2020, Zhao xiang et al. 2020).

Under the advocacy of the sustainable development practice known as “zero growth action plan for fertilizer use in China in 2020”, organic agriculture based on natural products such as humic acid and natural fertilizer has been implemented to improve soil quality. This practice is conducive to agricultural productivity and ecosystem health(Semida et al. 2019, Huang et al. 2020). The application of plant growth-promoting microorganisms to the soil is efficacious in organic agriculture. This approach compensates for the low efficiency of synthetic fertilizer, improves nutrient utilization efficiency and plant growth, reduces fertilizer investment by 50%, and causes no yield loss(Hayat et al. 2010, Good and Beatty 2011, Da Costa et al. 2013). However, organic fertilizer input may also create competition for nutrients between crops and soil microorganisms decomposing the fertilizer (Liu et al. 2009). I To balance this competition, integrated agricultural systems are nonetheless supplemented with chemical fertilizers to mitigate nutrient limitation (Harris 2009). I Plant probiotic enrichment in the form of organic substrates, such as bio-organic fertilizers, can help beneficial microorganisms thrive in the rhizosphere and improve their function (Yang et al. 2011, Mukta et al. 2017, Xiong et al. 2018, Bubici et al. 2019). Diacono and Montemurro (Diacono and Montemurro 2011) conducted over twenty long-term tests confirming that organic modifiers never lower crop yield. Zhang et al. (Zhang et al. 2016) showed that replacing 30% of the total nitrogen fertilizer (250 kg N/ha) with 3,000 kg/ha compost (equivalent to 60 kg N/ha) improved maize yield, N uptake, and soil fertility, and reduced N loss. Hence, there is a growing demand for bio-based organic fertilizers in agricultural production (Raja 2013).

Soil microorganisms regulate soil ecosystem functions and are indicators of soil quality (Sharma et al. 2010). They promote nutrient cycling and organic matter transformation, improve plant productivity, and control soil-borne diseases (Barrios 2007, Pieterse et al. 2016, Bakker et al. 2018). Microbial abundance, composition, and activity largely determine sustainable agricultural productivity (Van Der Heijden et al. 2008). The soil microbiome may be positively or negatively affected by soil disturbances and management practices that alter soil microbiome classification and function (Philippot et al. 2013, Cordovez et al. 2019). Long-term, large-scale chemical fertilizer use leads to the deterioration of soil quality, nutrient imbalances, reductions in soil microbial diversity, loss of microbial structural integrity, and a decrease in sustainable farmland productivity (Savci 2012). Bio-organic fertilizers contain beneficial microorganisms and organic components that directly or indirectly promote soil nutrient mobilization, have a positive impact on plant health, and improve crop yield (Tamreihao et al. 2016). Previous studies showed that bio-organic fertilizers stimulate specific microbiota associated with plant disease inhibition such as Pseudomonas, Streptomyces, Flavobacterium, and others (Mendes et al. 2011, Cha et al. 2016, Kwak et al. 2018). Therefore, strategies to stimulate the activities of these soil borne microbiota may be particularly effective in helping to inhibit plant diseases. Nevertheless, there are few reports on the changes that occur in the soil microbial diversity and root structure of Gansu Plateau summer vegetables in response to bio-organic fertilizer application.

In the present study, we conducted field trials on Baby Chinese cabbage under conventional fertilization, 30% chemical fertilizer reduction + 6,000 kg bio-organic fertilizer, 30% chemical fertilizer reduction + 9,000 kg bio-organic fertilizer, 40% chemical fertilizer reduction + 6,000 kg bio-organic fertilizer, and 40% chemical fertilizer reduction + 9,000 kg bio-organic fertilizer. We used directional sequencing of bacterial and fungal communities to analyze the response of the rhizosphere microbial community to the foregoing treatments. The aims of this study were to: 1) determine the changes in the physicochemical properties of the rhizosphere and the microbial community in response to various fertilizer treatments; 2) evaluate the impact of the different fertilizer treatments on Baby Chinese cabbage crop yield; 3) identify the correlations among rhizosphere microbial community composition, soil properties, and crop yield, 4) assess the benefits of combining bio-organic and chemical fertilizers, and 5) recommend the optimal amount of supplementary fertilizer required for Baby Chinese cabbage throughout its growth period. The results of this study will optimize crop productivity while minimizing the impact of chemical fertilization on the environment.

## 2. Materials and Methods

### 2.1 Experiment design and sampling

The experiment was conducted in Dachaigou Town (37°09′N, 102°99′E), Tianzhu County, Gansu Province between July 2019 and September 2020. This region has a continental plateau monsoon climate, altitude of ~2,630 m a.s.l., annual average temperature of −2 °C, annual average precipitation of 400–450 mm, and average annual sunshine hours of 2,500–2,700 h. The test site had a flat terrain and loam soil with medium, uniform fertility.

The Baby Chinese cabbage variety was “DeQin Golden Queen” produced by Beijing De Runcheng Agricultural Science and Technology Development Co. Ltd. (Beijing, China). The experiment was conducted over a 2-y period. In 2019, the Baby Chinese cabbage was planted on July 10 and harvested in September. In 2020, the Baby Chinese cabbage was planted in July and harvested in September. The fertilizers were urea (N ≥ 46%), calcium superphosphate (P_2_O_5_ ≥ 16%), potassium sulfate (K_2_O ≥ 52%) and bio-organic fertilizer (effective strains: Bacillus subtilis, Pseudomonas stutzeri, effective viable bacteria count ≥20 million/g, organic matter ≥ 40%). The calcium superphosphate and bio-organic fertilizer were applied as basic fertilizer. Thirty percent of the urea and potassium sulfate was applied as basal fertilizer, another 30% was applied at the rosette stage, and the remaining 40% was applied at early heading.

The experimental design comprised conventional fertilizer (CK), 30% chemical fertilizer reduction + 6,000 kg bio-organic fertilizer (T1), 30% chemical fertilizer reduction + 9,000 kg bio-organic fertilizer (T2), 40% chemical fertilizer reduction + 6,000 kg bio-organic fertilizer (T3), and 40% chemical fertilizer reduction + 9,000 kg bio-organic fertilizer (T4). Each plot was 0.008 ha in area and the cultivation mode was film-mulching ridge-furrow. The cultivation density was 59,550 plants/ha. The ridge and furrow widths were 50 cm and 40 cm, respectively, and the planting and row distances were 33 cm and 30 cm, respectively. Field management was consistent with local traditional cultivation. The fertilization rates under each treatment are listed in Table 1.

**Table 1.**
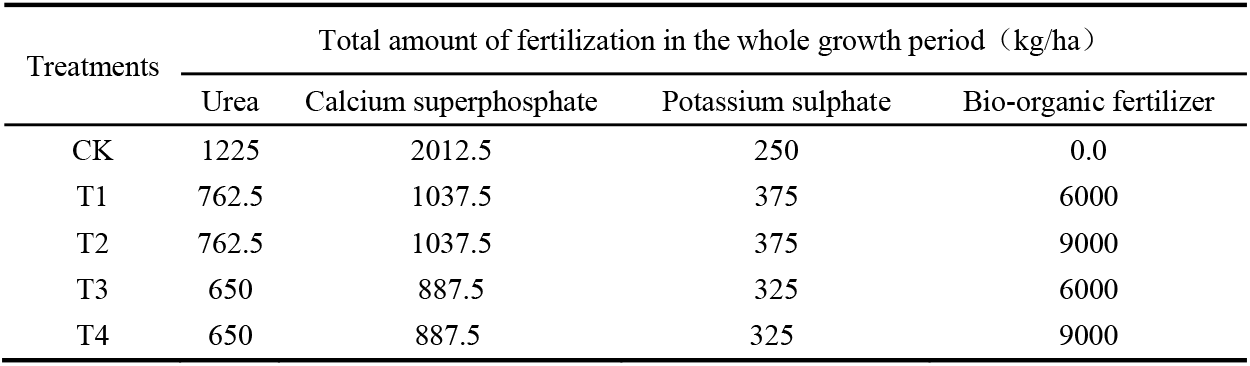
Fertilization rates for each treatment

| Treatments | Total amount of fertilization in the whole growth period (kg/ha) |  |  |  |
| --- | --- | --- | --- | --- |
|  | Urea | Calcium superphosphate | Potassium sulphate | Bio-organic fertilizer |
| CK | 1225 | 2012.5 | 250 | 0.0 |
| T1 | 762.5 | 1037.5 | 375 | 6000 |
| T2 | 762.5 | 1037.5 | 375 | 9000 |
| T3 | 650 | 887.5 | 325 | 6000 |
| T4 | 650 | 887.5 | 325 | 9000 |

Soil samples were collected on the Baby Chinese cabbage harvest date (September 11, 2020). For each plot, the edge effect was avoided. Thirty plants with the same growth were selected per plot. Surface soil was removed from 0 to 5 cm depth. Large soil clumps around the root systems were shaken off. The soil near the root systems was brushed with a brush, sealed in bags, placed in ice boxes, and taken to the laboratory. The soil samples were divided into two parts. One was air-dried and used in the physicochemical property determinations. The other was sieved through a 2-mm screen and stored at −80 °C until DNA extraction.

### 2.2 Analysis of soil physical and chemical properties

Soil physical and chemical properties were determined using the method of Lyu et al.(Lyu et al. 2020). Soil pH and EC were measured with a pH Meter (PHS-3E; Shanghai Jingke, Shanghai, China) and an EC Meter (DSJ-308Al Shanghai Jingke) (1:5 soil:water (w:v)), respectively. The soil organic matter content was measured by the potassium dichromate method. Total N, P, and K were determined by the Kjeldahl, molybdenum antimony colorimetry, and flame spectrophotometry (FP6410, Shanghai, China) methods, respectively.

### 2.3 DNA extraction and PCR amplification

Microbial genomic DNA extraction from soil samples was performed with the E.Z.N.A.® Soil DNA Kit (Omega Bio-tek, Norcross, GA, USA). DNA quality assays were performed on 1% agarose gels. DNA concentration and purity were determined with a NanoDrop 2000 UV-Vis spectrophotometer (Thermo Fisher Scientific, Wilmington, DE, USA) for determination. The V3-V4 region of bacterial 16S rRNA was amplified with primers s338F (5′-ACTCCTACGGGAGGCAGCAG-3′) and 806R (5′-GGACTACHVGGGTWTCTAAT-3′). The fungal ITS2 region was amplified with primers ITS1F (5′-CTTGGTCATTTAGAGGAAGTAA-3′) and ITS2R (5′-GCTGCGTTCTTCATCGATGC-3′). PCR was performed using an ABI GeneAmp® 9700 PCR thermocycler (Applied Biosystems, Foster City, CA, USA). The PCR amplification process was as follows: initial denaturation at 95°C for 3 min, denaturation at 95°C for 27 cycles of 30 s, annealing at 55°C for 30 s, extension at 72°C for 45 s, one-time extension at 72°C for 10 min, and termination at 4°C. A total volume of 20 μL of PCR mixture includes 4 μL of 5× TransStart FastPfu buffer, 2 μL of 2.5 mM dNTPs, 0.8 μL of 5 μM each of forward and reverse primers, 0.4 μL of TransStart FastPfu DNA polymerase, 10 ng of template DNA, and sufficient ddH2O. PCR Reactions were performed in triplicate. PCR products were electrophoresed on a 2% agarose gel, then purified using the AxyPrep DNA Gel Extraction Kit (Axygen Biosciences, Union City, CA, USA) and quantified using a Quantus™ fluorometer (Promega, Madison, WI, USA).

### 2.4 Illumina MiSeq sequencing

Purified amplicons were pooled in equimolar quantities and paired-end sequenced (2 × 300) according to standard protocols on an Illumina MiSeq platform (Illumina, San Diego, CA, USA) by Majorbio Bio-Pharm Technology Co. Ltd., Shanghai, China.

### 2.5 Processing of sequencing data

The raw 16S rRNA and ITS gene sequencing reads were demultiplexed, quality-filtered with Trimmomatic (https://github.com/usadellab/Trimmomatic), and merged by FLASH (https://sourceforge.net/projects/flashpage/files/) using the following criteria. (i) The 300-bp reads were truncated at any site with an average quality score < 20 over a 50-bp sliding window. Truncated reads < 50 bp and reads containing ambiguous characters were discarded. (ii) Only overlapping sequences > 10 bp were assembled. The maximum mismatch ratio of the overlap region was 0.2. Reads that could not be assembled were discarded. (iii) Samples were distinguished by their barcodes and primers and the sequence direction was adjusted. Exact barcode matching was used and two nucleotide mismatches were permitted in the primer matching.

The clean tags with sequence similarity greater than 97% were designated as an OTU using UPARSE v. 7.1 (http://drive5.com/uparse/) and then clustering was performed. The taxonomy of each OTU representative sequence (0.7 confidence threshold) was analyzed using the RDP classifier (http://rdp.cme.msu.edu/) against the fungal ITS (Unite 8.0) and bacterial 16S rRNA (Silva SSU128) databases.

### 2.6 Determination of yield

On September 16, 2019 and September 11, 2020, 30 plants were selected per plot and the yield per hectare was calculated based on the plot area and total yield.

### 2.7 Statistical analysis

Microbiological data analyses were conducted in R v. 3.5.2(Team 2018). The Shannon, Simpson, and Chao indices were calculated with QIIME (https://qiime2.org). A principal coordinate analysis (PCoA) was performed based on the Bray-Curtis distance and it evaluated the similarities of the microbial community composition among samples. A redundancy analysis (RDA) and Spearman’s rank correlation heat map analysis were used to study the relationships between soil physicochemical properties and soil microbial communities. Permutational multivariate ANOVA (PerMANOVA) was used to assess the effects of different fertilization regimes on soil microbial communities. PCoA, RDA, and PerMANOVA was implemented using the functions in the R vegan package (Oksanen et al. 2020). The Circos graph was plotted with Circos-0.67-7 (http://circos.ca/) (Yan et al. 2020). SPSS v. 21.0 (SPSS Inc., Chicago, IL, USA) was used to perform basic statistical tests such as one-way analysis of variance (ANOVA) and Pearson’s correlation analysis. Significant differences among treatments were indicated by P < 0.05 or other P values.

## 3. Result

### 3.1 Soil physicochemical properties under different fertilization regimes

The physicochemical properties of the soil significantly differed among fertilization treatments (Table 2). Compared with conventional fertilization (CK), the soil TN, TP, and TK increased with bio-organic fertilizer application rate. TN and TP were highest for the soil under T2 and were 23.08% and 39.76% greater than those under CK. Compared with CK, TP under T3 and T4 were 7.73% and 10.27% higher, and the differences were significant. The soil organic matter (SOM) content was lowest for CK and was significantly lower than the SOM under all other treatments. The SOM content increased sequentially from T1 to T4. All values were significantly higher than that of CK. The SOM content was highest under T4 and 48.08% greater than the SOM content under CK. The soil EC was highest under CK and was significantly higher than the soil EC under all other treatments. The soil EC under T1 through T4 were 37.49%, 29.28%, 24.36%, and 21.80% lower, respectively, than the soil EC under CK, and the differences were significant. The soil pH was highest under T2 and was 4.14% higher than the soil pH under CK. The difference was significant. The soil pH under all other treatments did not significantly differ from that under T2 but were significantly higher than that under CK.

**Table 2.** Physicochemical properties of soils under different fertilization conditions

| Treatments | TN<br>(g/kg) | TP<br>(g/kg) | TK<br>(g/kg) | SOM<br>(g/kg) | EC<br>(uS/cm) | pH |
| --- | --- | --- | --- | --- | --- | --- |
| CK | 0.39±0.02c | 0.83±0.02d | 16.17±0.30b | 33.38±0.58e | 507.67±5.21a | 7.25±0.07c |
| T1 | 0.48±0.01b | 0.94±0.06c | 17.17±0.44ab | 35.78±0.55d | 317.33±2.6d | 7.49±0.01ab |
| T2 | 0.54±0.03a | 1.16±0.03a | 17.42±0.30ab | 38.56±0.91c | 359.00±5.86c | 7.55±0.02a |
| T3 | 0.48±0.01b | 1.06±0.05b | 17.67±0.55a | 43.74±0.33b | 384.00±4.04b | 7.43±0.12ab |
| T4 | 0.50±0.01ab | 1.11±0.04ab | 17.83±0.51a | 49.43±0.77a | 397.00±5.87b | 7.48±0.08ab |

Values (mean ± standard deviation) followed by different lowercase letters within a column for the same factor indicate significant differences by Tukey test (P <0.05). TP, total phosphorus; TK, total potassium; TN, total nitrogen; pH, soil pH; EC, soil electrical conductivity; SOM, soil organic matter.

### 3.2 Changes in soil microbial α-diversity under different fertilization regimes

We measured the Shannon, Simpson, and Chao indices of bacteria (Fig. 1 A-C) and fungi (Fig. 1 D-F) to evaluate the α-diversity of the soil microbial community in the Baby Chinese cabbage rhizosphere under different fertilization treatments. The Shannon index is directly proportional to the diversity while the Simpson index is inversely proportional to it. The Chao index is directly proportional to the microbial abundance. Figure 1-A shows that after the various fertilization treatments, the Shannon index was 0.63% and 0.78% higher for the soil bacteria under T1 and T3, respectively, than it was for those under CK and the differences were significant. There was no significant difference between T4 and CK in terms of the Shannon index. However, the Shannon index was 1.10% lower under T2 than CK and the difference was significant. There were no significant differences among fertilization treatments in terms of their bacterial Simpson indices (Fig. 1-B). The bacterial Chao index was highest under T4 and was 34.20% higher than that under CK. However, there were no significant differences among the other treatments and CK in terms of the bacterial Chao index (Fig. 1-C). The fungal Shannon indices under T1 through T4 were 10.66, 14.42, 8.46, and 14.73% higher, respectively, than that under CK, and the differences were significant (Fig. 1-D). The fungal Simpson indices under T1 through T4 were 44.53, 53.13, 34.38 and 50.78% lower, respectively, than that under CK, and the differences were significant (Fig. 1-E). The fungal Chao indices under T2 and T4 were 23.40% and 23.49% higher, respectively, than that under CK, and the differences were significant (Fig. 1-F).

**Fig. 1.**
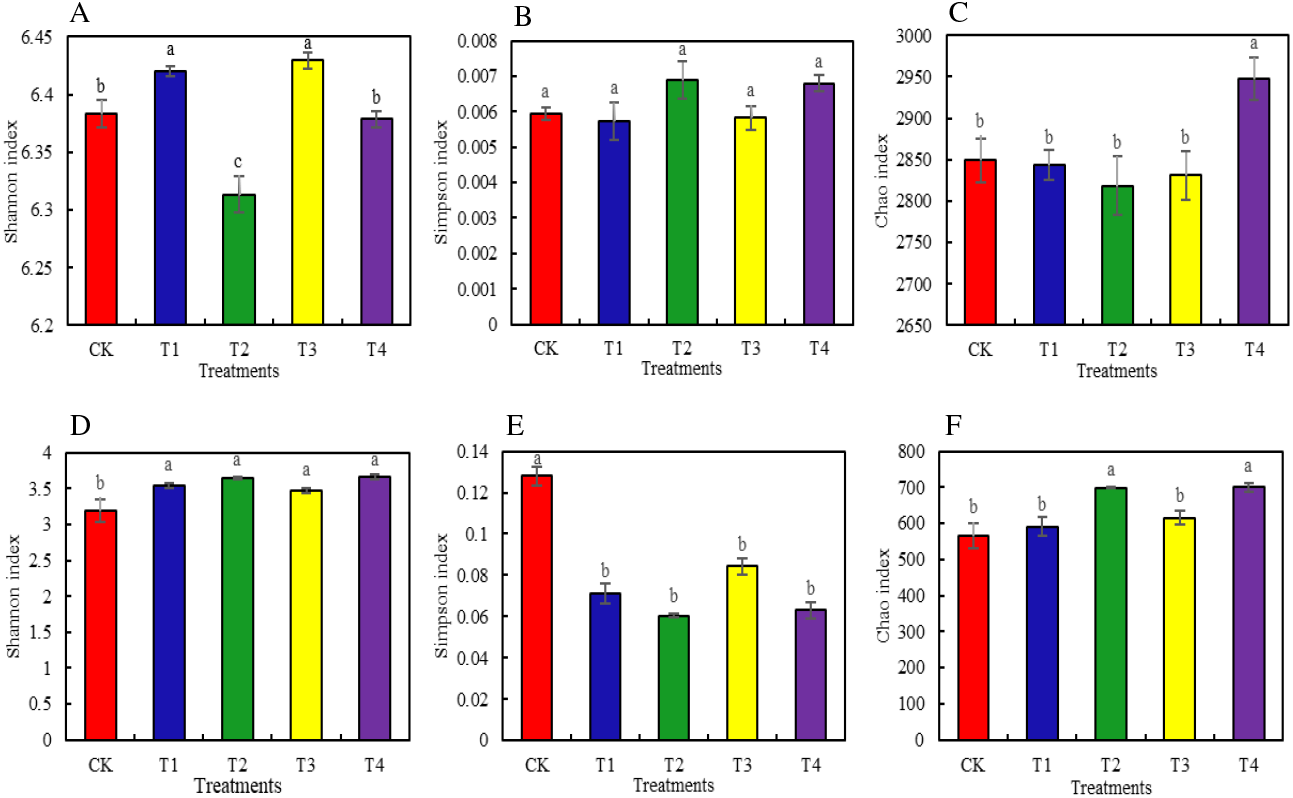
The α-diversity of bacteria (A-C) and fungi (D-F) in the root soil of Baby Chinese cabbage under different fertilization treatments. Different letters over the plots indicate significant differences (P < 0.05).

### 3.3 Changes in soil microbial α-diversity under different fertilization regimes

We applied PCoA based on the Bray-Curtis algorithm to analyze overall structural changes in the bacterial and fungal microbial communities and determine the impact of reducing chemical fertilizers and adding bio-organic fertilizers on their structure. The fertilization regimes significantly altered soil bacterial and fungal community structure in the Baby Chinese cabbage rhizospheres. The PCoA assigned the bacterial communities to two groups, namely, [CK, T1, and T3] and [T2 and T4] (Fig. 2-A). The T2 and T4 bacterial community structures were clustered together and were remote from those of the other treatment groups (Fig. 2-A). Hence, the soil bacterial community structure significantly changed with increasing bio-organic fertilizer application rate. The PCoA assigned the fungal communities to three groups, namely, [CK], [T1 and T3], and [T2 and T4] (Fig. 2-B). The T2 and T4 fungal community structures were clustered together and were remote from that of CK (Fig. 2-B). There was a certain distance between T1 and T3 but both were nonetheless in the same quadrant. Thus, the change in the fungal community structure of the Baby Chinese cabbage rhizosphere depended on whether biological organic fertilizer was used and the quantity added. PERMANOVA revealed that each fertilization treatment significantly affected the bacterial (*F* = 1.64, *P* = 0.01) and fungal (*F* = 6.66, *P* = 0.01) community structures.

**Fig. 2.**
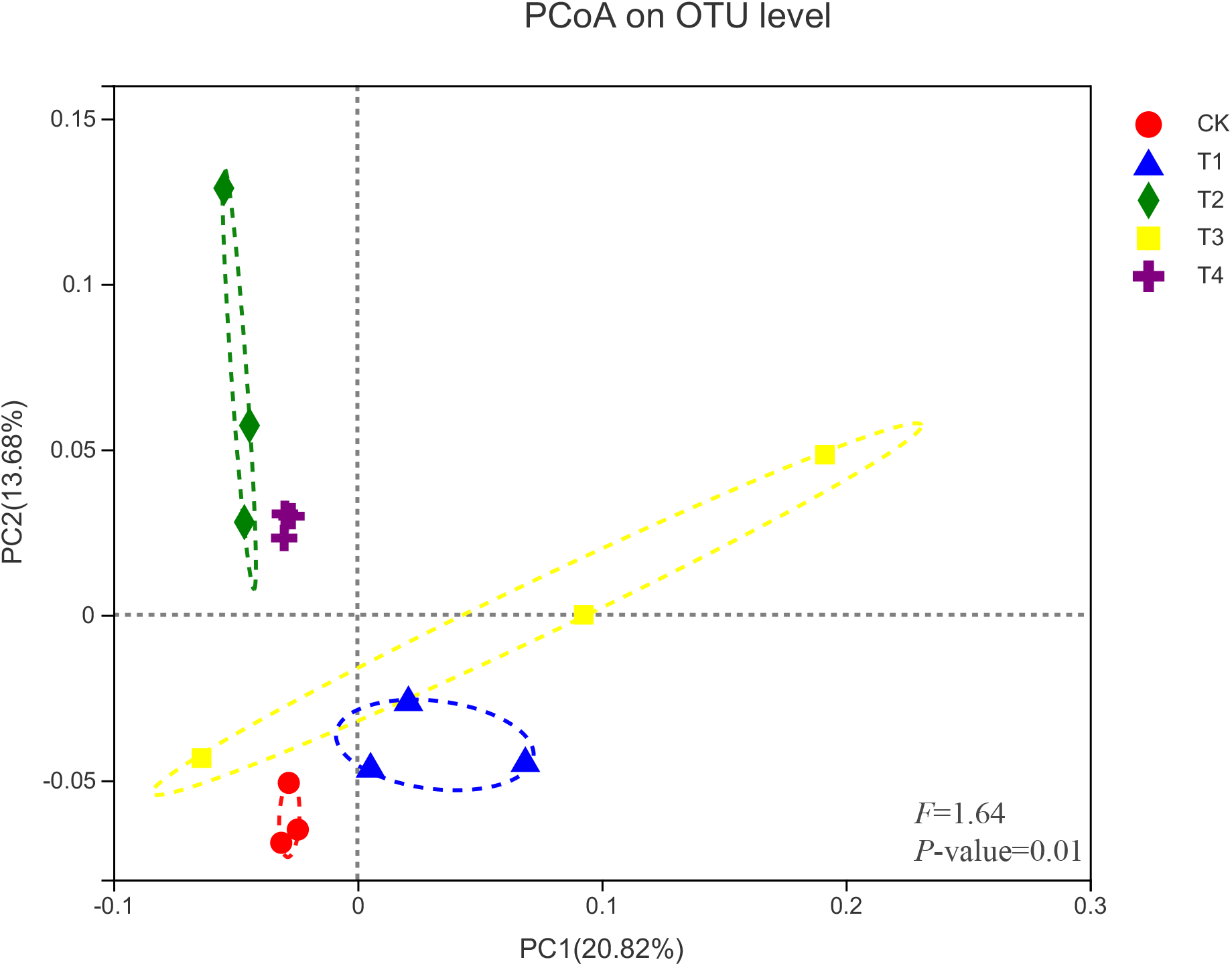
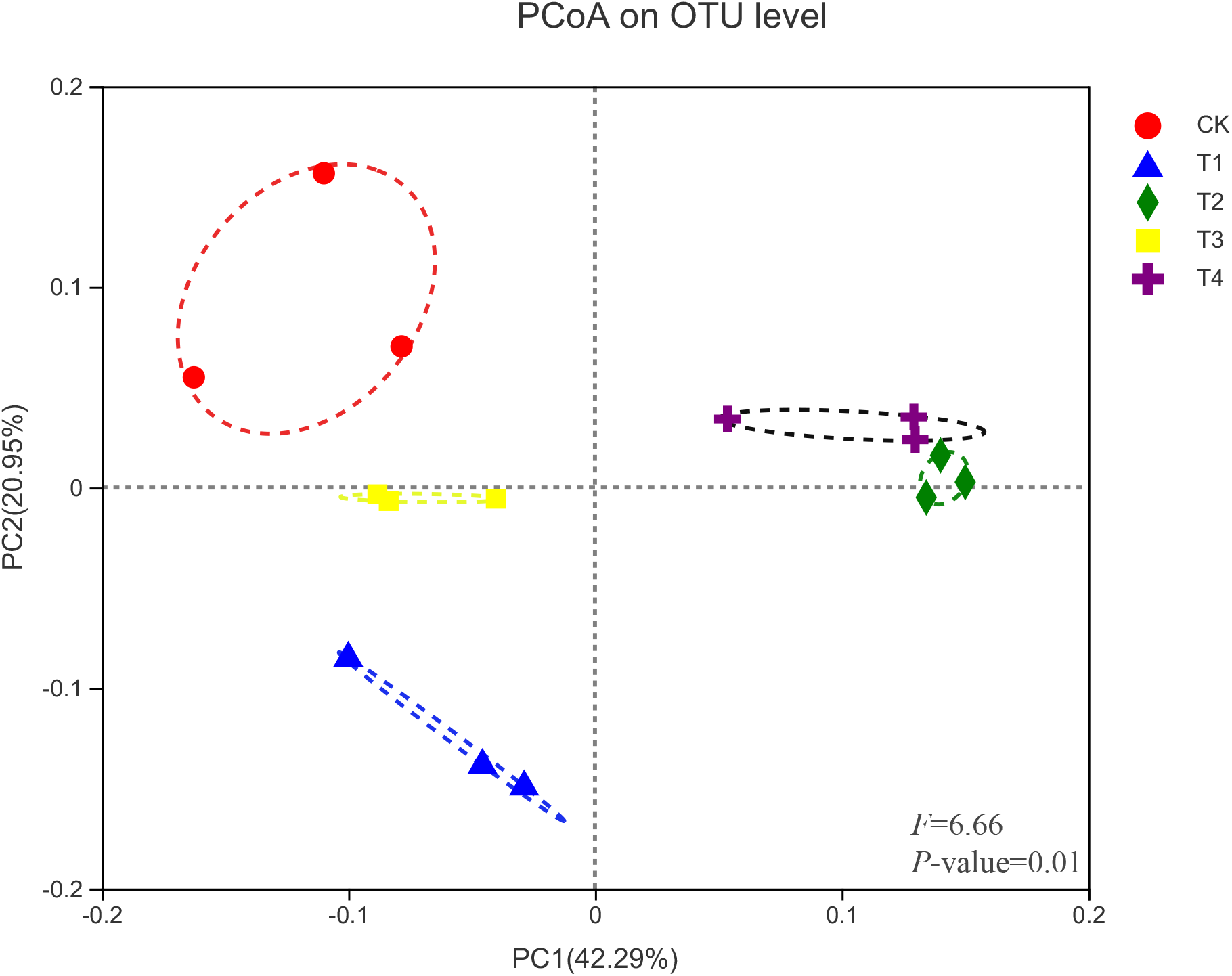
Principal coordinate analysis (PCoA) of the bacteria (A) and fungi (B) communities based on the Bray–Curtis distances.

### 3.4 Associations between soil microbial communities and environmental factors

A redundancy analysis (RDA) was used to evaluate the influences of environmental factors on the soil microbial community composition in Baby Chinese cabbage rhizosphere (Fig. 3). The changes in the soil properties in response to the different fertilization treatments strongly affected the soil bacterial and fungal community structures. The axes explained that the variation in the soil bacterial (Fig. 3-A) and fungal (Fig. 3-C) communities were 19.63 and 81.56%, respectively. The first axis in Fig. 3-A was positively correlated with soil TN, TK, TP, pH, and SOM and negatively correlated with soil EC. The second axis was positively correlated with soil EC, TP, and pH and negatively correlated with soil TN, TK, and SOM. We also used Spearman’s rank correlation to evaluate the relationships among the abundant bacterial phyla and soil physicochemistry (Fig. 3-B). Bacteroidota were significantly positively correlated with soil TN and TK, SAR324_cladeMarine_group_B and Firmicutes were significantly negatively correlated with soil EC, Acidobacteriota were significantly negatively correlated with soil TN and pH while Methylomirabilota and Latescibacterota were significantly negatively correlated with SOM. The first axis in Fig. 3-B was positively correlated with soil TN, TK, TP, pH, and SOM and negatively correlated with soil EC. The second axis was positively correlated with soil EC and SOM and negatively correlated with soil TN, TK, TP, and pH. Spearman’s rank correlation between the soil fungal community and physicochemistry disclosed that Zoopagomycota, unclassified_k__Fungi, and Mortierellomycota were significantly positively correlated with soil TP, SOM, and TN, respectively, and Ascomycota were significantly positively correlated with soil TK, SOM, and TN and very significantly positively correlated with soil pH. Mortierellomycota, Kickxellomycota, and Blastocladiomycota were significantly negatively correlated with soil EC while Olpidiomycota were significantly negatively correlated with SOM and soil pH.

**Fig. 3.**
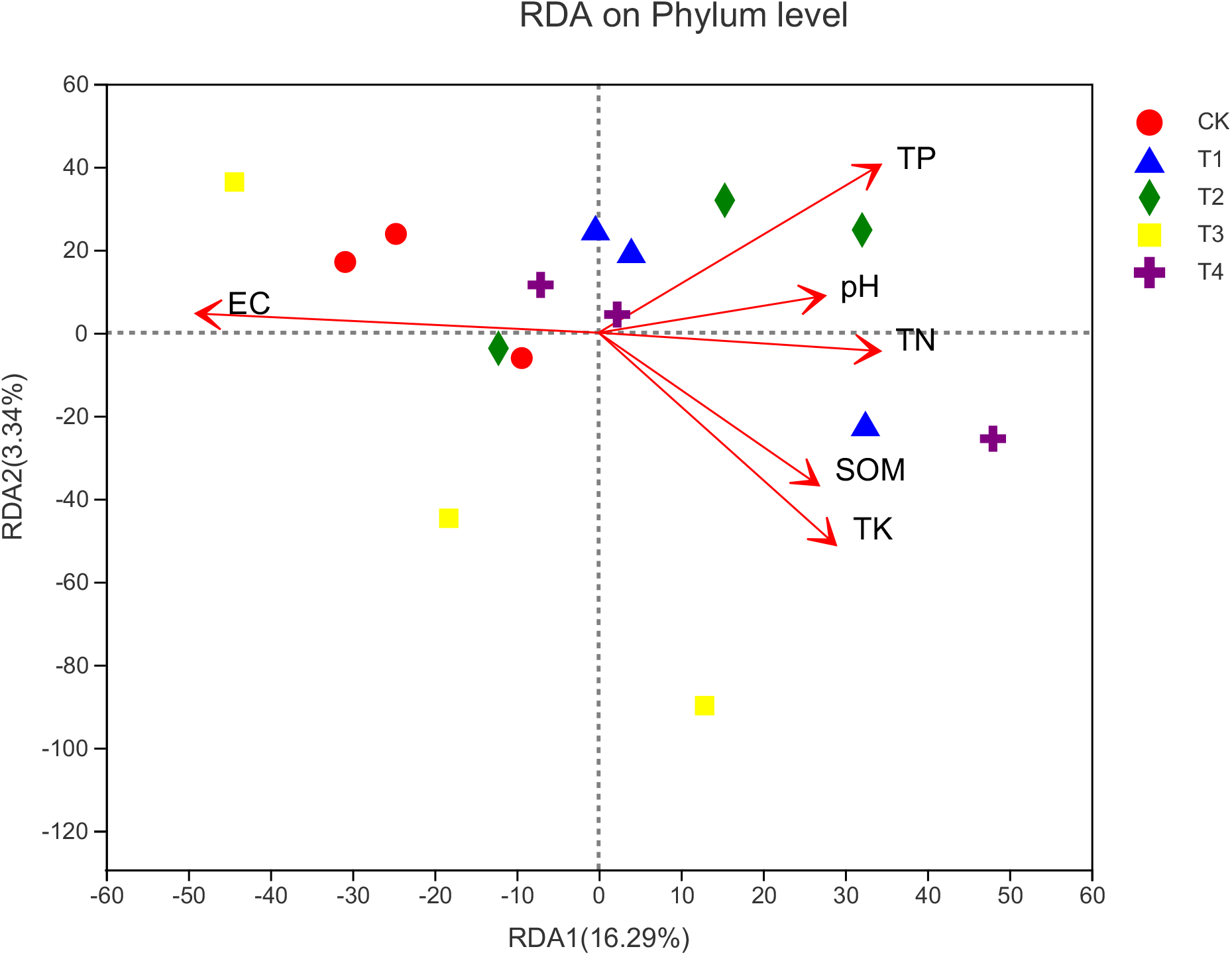
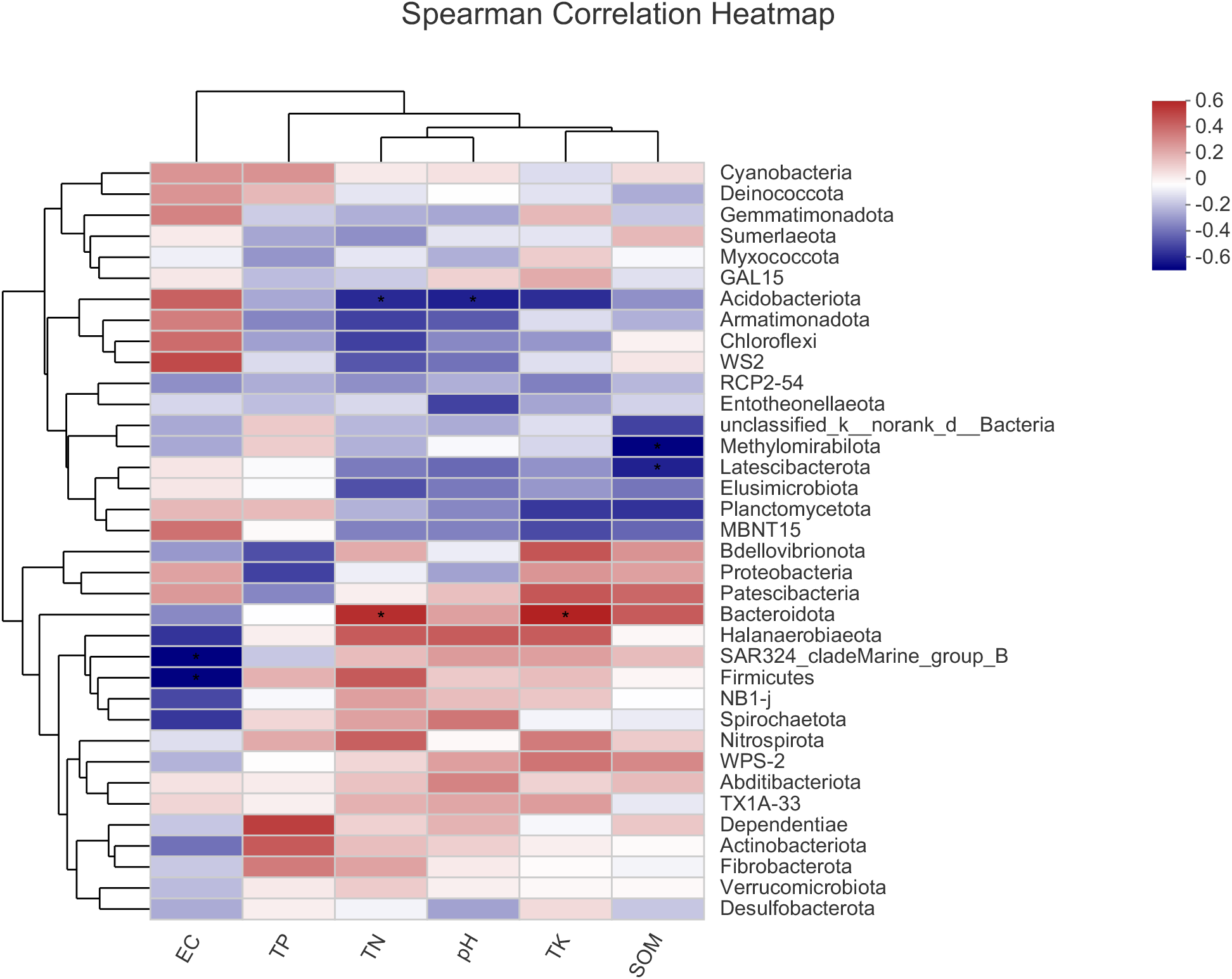
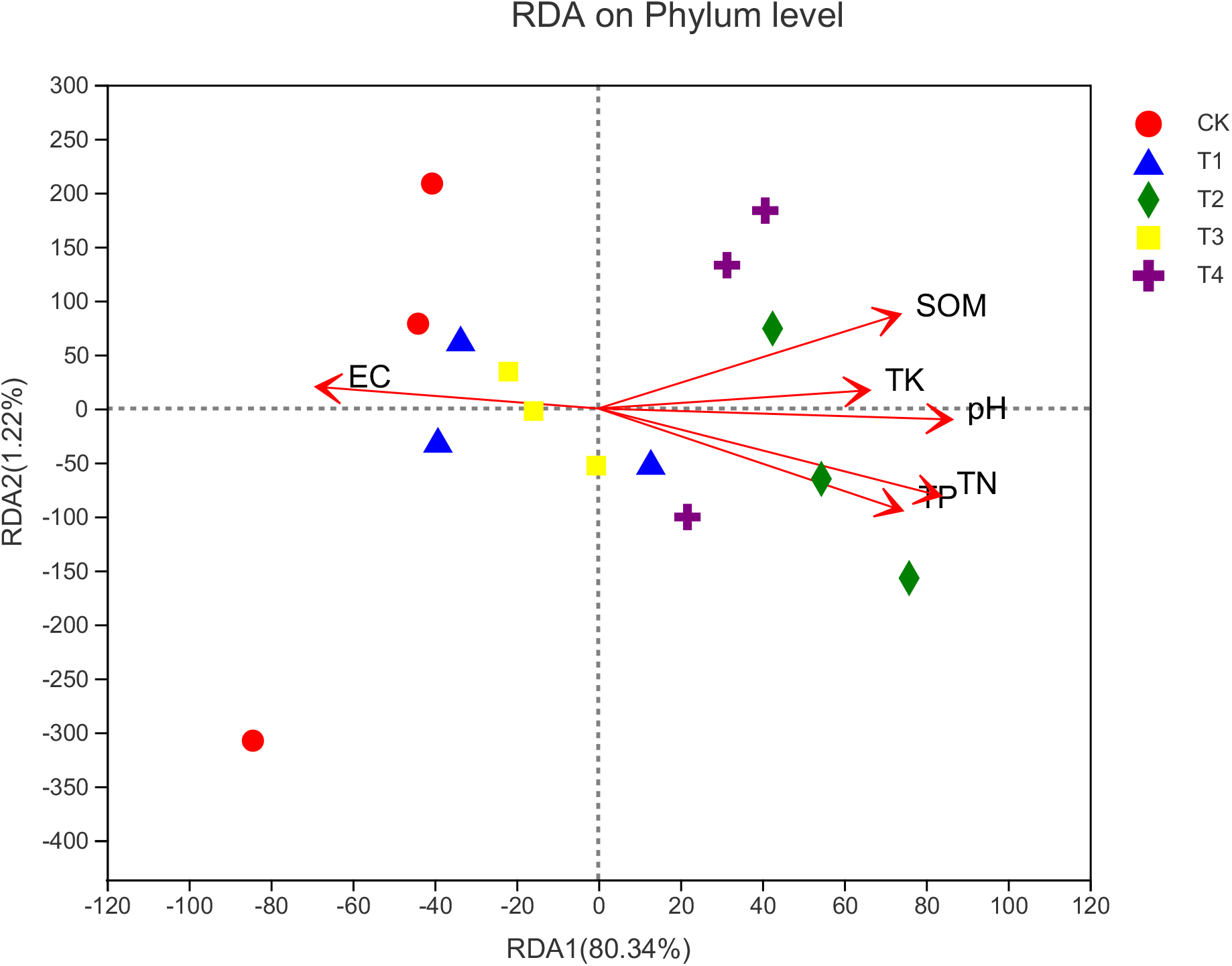
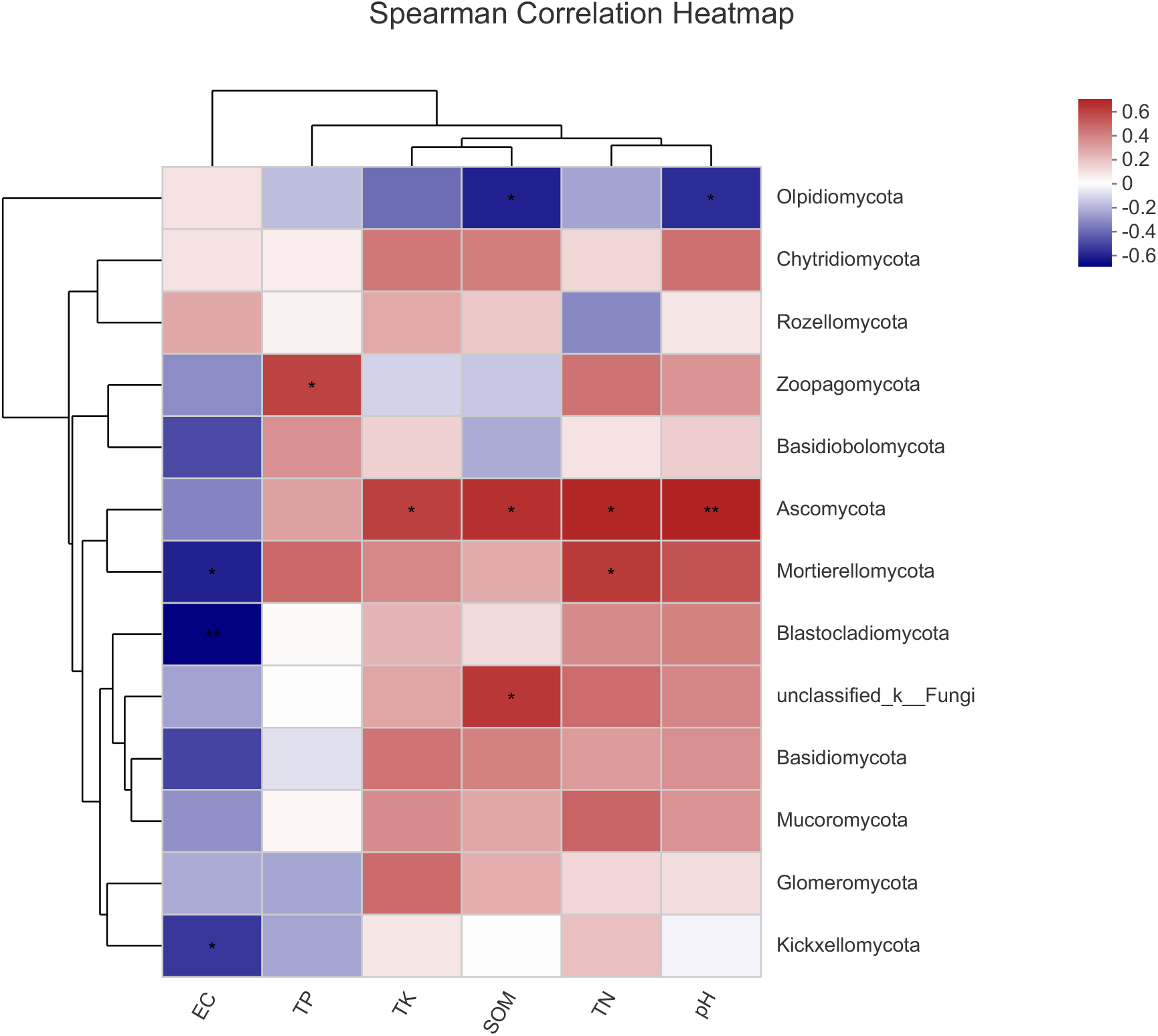
Redundancy analysis (RDA) and Spesrman’s rank correlation heatmap (P < 0.05 *, P < 0.01**), used to study the correlation between the microbial communities of bacteria (A-B) and fungi (C-D) and soil physical and chemical properties. TP, total phosphorus; TK, total potassium; TN, total nitrogen; pH, soil pH; EC, soil electrical conductivity; SOM, soil organic matter.

### 3.5 Relative abundance of major bacterial and fungal taxa

The main bacterial phyla were Actinobacteriota, Proteobacteria, Chloroflexi, and Acidobacteriota (Fig. 4-A) and their average relative abundances were 35.14, 19.83, 16.39, and 10.56%, respectively. The subdominant bacterial phyla were Gemmatimonadota, Bacteroidota, Myxococcota, and Firmicutes. T1 through T4 had higher actinobacteria richness than that of CK. T2 and T4 had the highest and second highest Actinobacteriota abundance. Actinobacteriota abundance was higher under T1 than T3. T3 and T4 had higher Proteobacteria abundance than CK. However, Proteobacteria abundance did not significantly differ between T1 and T2. Compared with CK, Chloroflexi, Acidobacteriota, and Gemmatimonadota had higher abundance while Bacteroidota and Firmicutes had lower abundance under T1 through T4. The main fungal taxa were Ascomycota, Olpidiomycota, Mortierellomycota, Basidiomycota, and unclassified_k__Fungi (Fig. 4-B). T1 through T4 had higher Ascomycota abundance than CK. T2 had the highest Ascomycota abundance but relatively low Olpidiomycota abundance. T1 through T4 had relatively high Mortierellomycota abundance.

**Fig. 4.**
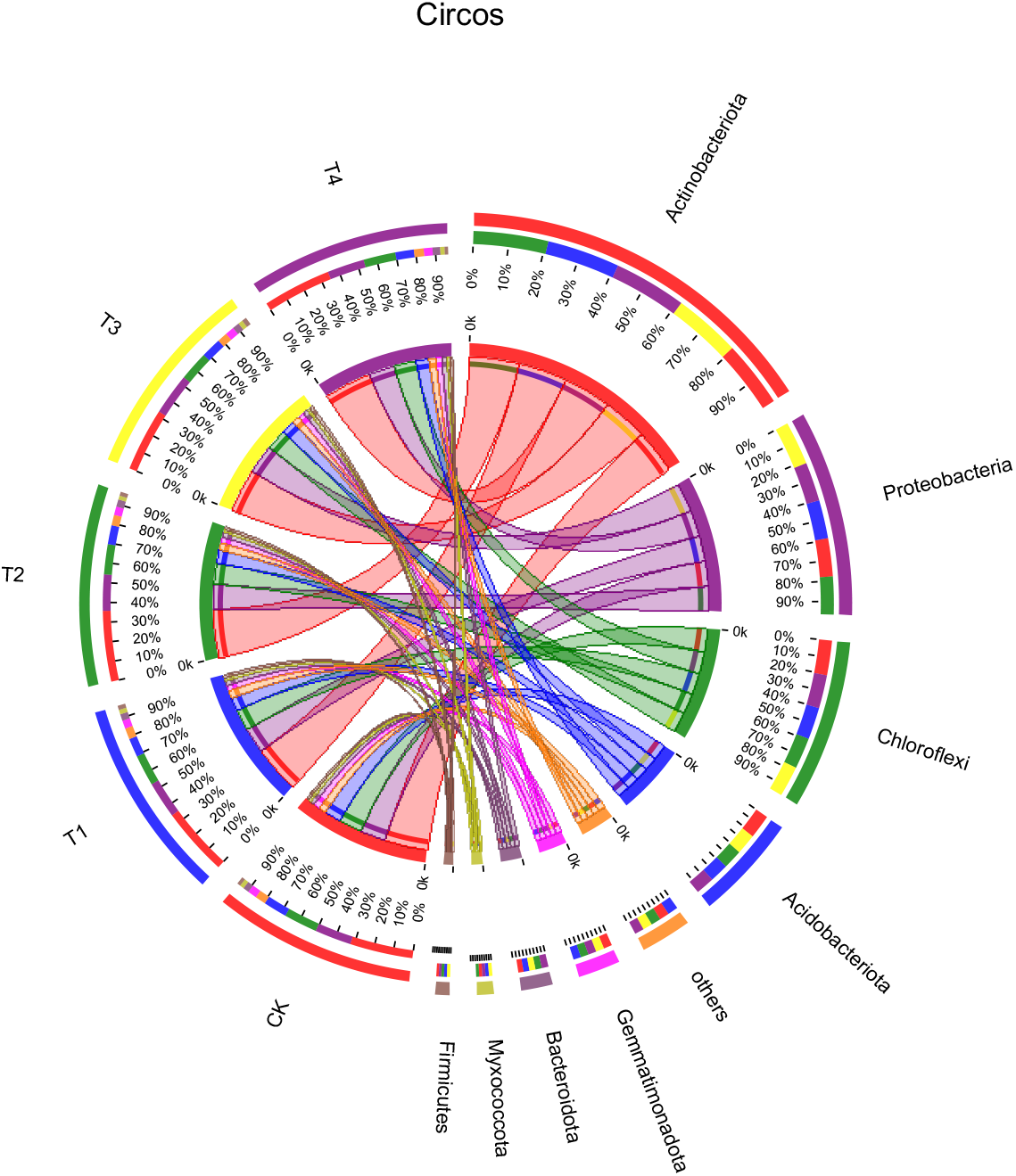
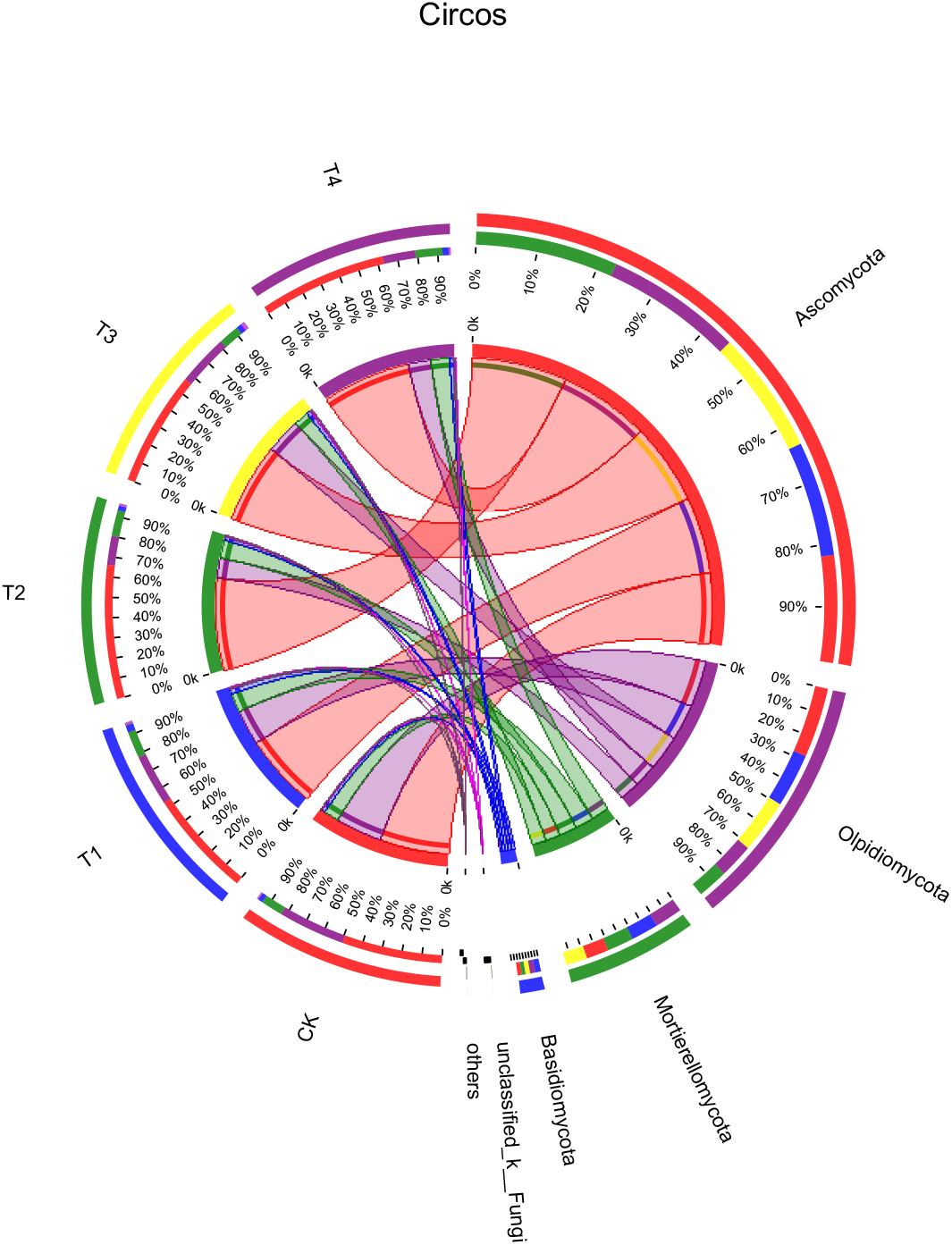
The relative abundance of major taxonomic groups at the phylum level (Others incorporated<0.01) for bacteria (A) and fungi (B). The data were visualized by Circos. The width of the bars from each phylum indicated the relative abundance of the phylum.

### 3.6 Effect of different fertilization regimes on Baby Chinese cabbage yield

We measured Baby Chinese cabbage yield in 2019 and 2020 and calculated the average for both years. Compared with CK, T1 through T4 significantly increased Baby Chinese cabbage yield by 6.39, 8.89, 3.94, and 5.14%, respectively. The yield was highest under T2 but did not significantly differ from that under T1 (Fig. 5).

**Fig. 5.**
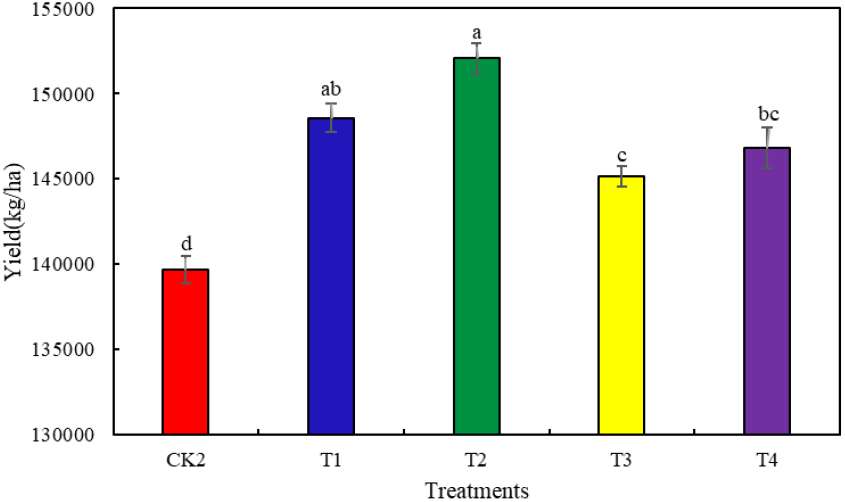
Average yield of Baby Chinese cabbage under different fertilization systems in 2019–2020. Different letters over the plots indicate significant differences (P < 0.05).

## 4. Discussion

Widespread use of chemical fertilizers has led to losses of soil quality and organic carbon content (Kumar et al. 2017, Kumar et al. 2018). Therefore, fertilization must be optimized to ensure sustainable food production. Compared with conventional soil cultivation, organic soil cultivation enhances microbial functional diversity. Long-term application of organic fertilizers or organic-inorganic compound fertilizers may increase microbial biomass and enzyme activity and improve SOM quantity and quality (Bending et al. 2004). In our study, compared with conventional fertilization, all four chemical fertilizer reduction and combined with bio-organic fertilizer application treatments significantly increased soil TP, TN, TK, pH, and SOM. Furthermore, both TK and SOM increased with bioorganic fertilizer application rate (Table 2). Therefore, the combined application of organic and chemical fertilizers increases soil fertility and supports sustainable productivity more effectively than chemical fertilizer application alone (Liu et al. 2014, Jiang et al. 2017), The addition of Bacillus subtilis and Pseudomonas stutzeri to the organic fertilizer further increased soil extracellular enzyme activity and nutrient content (Li et al. 2021). TP, TN, and pH were at their maxima under T2 (Table 2). For this reason, even moderate reductions in chemical fertilizer application rate followed by bio-organic fertilizer treatment effectively increase soil nutrient content and improve soil quality. By contrast, excessive reductions in chemical fertilizer application may decrease soil nutrient levels even when sufficient bio-organic fertilizer is used.

Several studies evaluated the effects of fertilization on soil microbes. Organic fertilizers, such as manure might have a positive effect on the soil microbial community. Compared with chemical or no fertilization, organic fertilization improves the resistance of the soil microbial community to perturbations (Francioli et al. 2016, Cui et al. 2018, Legrand et al. 2018). Our results showed that compared with conventional fertilization, T1 and T3 significantly increased bacterial diversity and T4 significantly increased bacterial richness (Fig. 1A-C). These findings were consistent with those of prior studies indicating that organic fertilizer addition increases bacterial community diversity (Chaudhry et al. 2012, Jannoura et al. 2013, Xiong et al. 2017). Compared with conventional fertilization, organic fertilization had a stronger effect on soil fungal communities (Cwalina-Ambroziak and Bowszys 2009, Wei et al. 2017). All treatments with bio-organic fertilizer increased soil fungal community diversity and richness. However, T2 and T4 were the most effective (Fig. 1D-F).That is, the effective enhancement of soil microbial diversity and richness may be a combined effect of Bioorganic and chemical fertilizer applications (Ding et al. 2018).

The soil microbial community structure significantly differed between the bio-organic fertilizer and other treatments. The PCoA showed that the bacterial and fungal community structures under T2 and T4 and those under T1 and T3 were similar. In both cases, they were separated by conventional fertilization (CK) (Fig. 2A-B). These discoveries were consistent with previous observations that soil microbial community structures differ in response to organic and inorganic fertilizers (Marschner et al. 2003, Zhang et al. 2012). Moreover, fertilization regimes more strongly influence fungal than bacterial community structure (Wang et al. 2018b, Zhang et al. 2018).

A combination of PCoA and RDA demonstrated that fertilization may modulate the impact of environmental factors on fungal community structure. The variance between fungal community structure and environmental factors was 81.56% (Fig. 3-C). By contrast, the variance between bacterial community structure and environmental factors was only 19.63% (Fig. 3-A). Fertilizer application increases the abundance of oligotrophic organisms such as Bacteroidota which are significantly positively correlated with soil TN and TK (Fig. 3-B). Oligotrophic organisms tend towards K-selection (Berg and Smalla 2009). A high relative abundance of K-strategists promotes microbial community resistance and stability (Simonin et al. 2017). Soil pH is positively correlated with most bacterial communities but negatively correlated with Acidobacteriota (Fig. 3-B). In the present study, the soil pH was near neutrality and organic fertilizer application prevented soil acidification. In this manner, it buffered the potential effects of soil acidity on bacterial richness and diversity (Zhang et al. 2017). However, fungal communities are comparatively more strongly affected by the type of fertilization system. Fungal richness is positively correlated with soil N availability (Hu et al. 2017). Ascomycota are the most abundant fungi in the Baby Chinese cabbage rhizosphere and are significantly positively correlated with TN (Fig. 3-D). Hence, the fungal community is closely related to the soil nitrogen fertilizer application rate. SOM also plays an important role in fungal community composition (Yao et al. 2017, Dai et al. 2018). Most dominant fungal phyla such as Ascomycota, Zoopagomycota, unclassifìed_k__Fungi, and Mortierellomycota are positively correlated with SOM. Increases in EC are the result of increases in salinity caused by continuous soil cropping. Soil salinity has negative effects on soil microbial communities (Shen et al. 2016). The application of bio-organic fertilizer can reduce EC. Most dominant fungal phyla are negatively correlated with this parameter (Fig. 3-D).

In all fertilization treatments, Actinobacteria, Proteobacteria, Chloroflexi, and Acidobacteria were the dominant bacterial phyla (Fig. 4-A). This observation was consistent with those of previous studies (Guo et al. 2018, Huang et al. 2019). Actinobacteriota comprise a group of co-nutrients suitable for plant growth in high-C environments (Fierer et al. 2012), Bio-organic fertilizers promote Actinobacteriota proliferation because they create nutrient- and carbon-rich environments and their abundance significantly increased in response to bio-organic fertilizer treatment and was highest under T2. Composting is beneficial to the growth of Proteobacteria and Firmicutes (Martínez-García et al. 2018). In this study, bio-organic fertilizer treatment also increased Proteobacteria and Firmicutes abundance. Chloroflexi include numerous acid-producing bacteria that anaerobically digest food waste and produce methane gas (Wang et al. 2018a). Long-term chemical fertilizer application may acidify the soil (Guo et al. 2010), thereby reducing soil microbial diversity and community structure (Zhang et al. 2015). In the present study, Chloroflexi abundance was higher under the conventional fertilization treatment. The addition of bio-organic fertilizer increased the soil pH (Table 2) and reduced Chloroflexi abundance. Acidobacteria are slow-growing oligotrophs that flourish under low-nutrient conditions. Their growth is inhibited by N input (Trivedi et al. 2017). Bio-organic fertilizer addition increased soil nutrient levels and inhibited Acidobacteria growth. Nevertheless, further research is required to elucidate the mechanism(s) by which bio-organic fertilizer addition increases bacterial abundance.

Ascomycota were the dominant fungi in the present study (Fig. 4-B). These saprophytes can thrive in arid environments, have strong environmental adaptability, degrade organic substrates, and are major decomposers of SOM containing cellulose, lignin, and pectin (Yelle et al. 2008). The combination of bio-organic fertilizer plus chemical fertilizer reduction increased Ascomycota abundance. Ascomycota richness was highest under T2. Mortierellomycota are saprophytic and ubiquitous. They can dissolve phosphorus, increase crop yield, and form symbiotic relationships with plants (Fröhlich-Nowoisky et al. 2015, Grządziel and Gałązka 2019). In the present study, Mortierellomycota abundance increased in response to bio-organic fertilizer treatment and was significantly positively correlated with TP, SOM, and TN (Fig. 3-D). Therefore, improvements in the soil nutrients in Baby Chinese cabbage rhizosphere may be associated with higher relative Mortierellomycota abundance.

The production and integration of new bio-organic fertilizers with beneficial microorganisms and mature compost is an important way to increase crop yields and/or strengthen the control of soil-borne diseases(Zheng et al. 2020, Chen et al. 2021). It is feasible but difficult to reduce fertilizer application without causing a loss of productivity (Da Costa et al. 2013). The results of this experiment confirmed our hypothesis that bio-organic fertilizer significantly increased Baby Chinese cabbage yield compared with chemical fertilizer. T2 (30% chemical fertilizer reduction + 9,000 kg bio-organic fertilizer) realized the highest crop yield of all treatments. Ye et al.(Ye et al. 2020) showed that a combination of chemical fertilizer reduction and bio-organic fertilizer application could improve crop yield and quality.

These results provide a foundation for understanding how the addition of bioorganic fertilizers caused soil physicochemical improvement and then caused changes in soil microbial diversity and richness, which in turn enhanced Baby Chinese cabbage yield (Fig. 6).

**Fig. 6.**
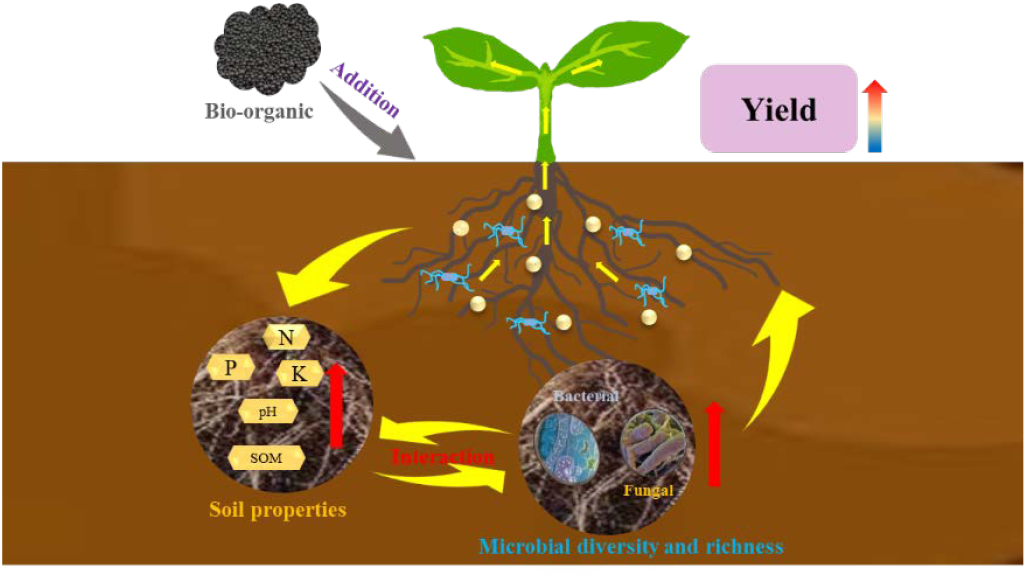
A model of soil physical and chemical properties, microbial community and yield changes after adding bio-organic fertilizer.

## 5. Conclusion

Overall, the present study examined the effects of various fertilization regimes on soil microbial diversity and community structure in the rhizosphere of Baby Chinese cabbage grown on the Gansu Plateau. The potential value of bio-organic fertilization in Baby Chinese cabbage cultivation was assessed in terms of microbial structure and crop yield. Reduction of chemical fertilizers and addition of bio-organic fertilizers improved rhizosphere physicochemistry, markedly altered the soil bacterial and fungal communities, increased their diversity and abundance, and improved the soil microenvironment. Compared with conventional fertilization, the combination of bio-organic fertilizer and reduced chemical fertilizer created a richer, more diverse microenvironment and realized higher Baby Chinese cabbage yield. Hence, this novel approach is relatively more efficient and desirable in terms of sustainable and high crop yield. Future research should investigate the changes that occur in the microbial profile of the rhizosphere and the effects of these microorganisms on the yield and quality of Baby Chinese cabbage subjected to long-term bio-organic fertilization.

## Data Availability Statement

The datasets generated during and/or analysed during the current study are available from the corresponding author on reasonable request. Sequencing data are stored in the NCBI database, access link: https://identifiers.org/ncbi/insdc.sra:SRP359012

## Acknowledgements

This research was funded by the Education science and technology innovation project of Gansu Province (GSSYLXM-02), the Special project of central government guiding local science and technology development (ZCYD-2021-07), Gansu people’s livelihood science and technology project (20CX9NA099), the Fuxi Young Talents Fund of Gansu Agricultural University (GAUfx-04Y03), Gansu Top Leading Talent Plan (GSBJLJ-2021-14).

This article thanks the experimental field provided by Tianzhu experimental station, as well as the editors and several reviewers for their valuable comments on this article.

## References

Ajeng, A. A., R. Abdullah, T. C. Ling, S. Ismail, B. F. Lau, H. C. Ong, K. W. Chew, P. L. Show, and J.-S. Chang. 2020. Bioformulation of biochar as a potential inoculant carrier for sustainable agriculture. Environmental Technology & Innovation:101168.

Bakker, P. A., C. M. Pieterse, R. de Jonge, and R. L. Berendsen. 2018. The soil-borne legacy. Cell 172:1178–1180.

Barrios, E. 2007. Soil biota, ecosystem services and land productivity. Ecological economics 64:269–285.

Bending, G. D., M. K. Turner, F. Rayns, M.-C. Marx, and M. Wood. 2004. Microbial and biochemical soil quality indicators and their potential for differentiating areas under contrasting agricultural management regimes. Soil Biology and Biochemistry 36:1785–1792.

Berg, G., and K. Smalla. 2009. Plant species and soil type cooperatively shape the structure and function of microbial communities in the rhizosphere. FEMS microbiology ecology 68:1–13.

Bubici, G., M. Kaushal, M. I. Prigigallo, C. Gómez-Lama Cabanás, and J. Mercado-Blanco. 2019. Biological control agents against Fusarium wilt of banana. Frontiers in Microbiology 10:616.

Cha, J.-Y., S. Han, H.-J. Hong, H. Cho, D. Kim, Y. Kwon, S.-K. Kwon, M. Crüsemann, Y. B. Lee, and J. F. Kim. 2016. Microbial and biochemical basis of a Fusarium wilt-suppressive soil. The ISME journal 10:119–129.

Chaudhry, V., A. Rehman, A. Mishra, P. S. Chauhan, and C. S. Nautiyal. 2012. Changes in bacterial community structure of agricultural land due to long-term organic and chemical amendments. Microbial ecology 64:450–460.

Chen, Y., X. Li, S. Li, and Y. Xu. 2021. Novel-integrated process for production of bio-organic fertilizer via swine manure composting. Environmental Engineering Research 26:22–33.

Cordovez, V., F. Dini-Andreote, V. J. Carrión, and J. M. Raaijmakers. 2019. Ecology and evolution of plant microbiomes. Annual review of microbiology 73:69–88.

Cui, X., Y. Zhang, J. Gao, F. Peng, and P. Gao. 2018. Long-term combined application of manure and chemical fertilizer sustained higher nutrient status and rhizospheric bacterial diversity in reddish paddy soil of Central South China. Scientific reports 8:1–11.

Cwalina-Ambroziak, B., and T. Bowszys. 2009. Changes in fungal communities in organically fertilized soil. Plant Soil Environ 55:25–32.

Da Costa, P. B., A. Beneduzi, R. de Souza, R. Schoenfeld, L. K. Vargas, and L. M. Passaglia. 2013. The effects of different fertilization conditions on bacterial plant growth promoting traits: guidelines for directed bacterial prospection and testing. Plant and soil 368:267–280.

Dai, Z., A. Enders, J. L. Rodrigues, K. L. Hanley, P. C. Brookes, J. Xu, and J. Lehmann. 2018. Soil fungal taxonomic and functional community composition as affected by biochar properties. Soil Biology and Biochemistry 126:159–167.

Diacono, M., and F. Montemurro. 2011. Long-term effects of organic amendments on soil fertility. Sustainable agriculture volume 2:761–786.

Ding, L.-J., J.-Q. Su, G.-X. Sun, J.-S. Wu, and W.-X. Wei. 2018. Increased microbial functional diversity under long-term organic and integrated fertilization in a paddy soil. Applied microbiology and biotechnology 102:1969–1982.

Fierer, N., C. L. Lauber, K. S. Ramirez, J. Zaneveld, M. A. Bradford, and R. Knight. 2012. Comparative metagenomic, phylogenetic and physiological analyses of soil microbial communities across nitrogen gradients. The ISME journal 6:1007–1017.

Francioli, D., E. Schulz, G. Lentendu, T. Wubet, F. Buscot, and T. Reitz. 2016. Mineral vs. organic amendments: microbial community structure, activity and abundance of agriculturally relevant microbes are driven by long-term fertilization strategies. Frontiers in Microbiology 7:1446.

Fröhlich-Nowoisky, J., T. C. Hill, B. G. Pummer, P. Yordanova, G. D. Franc, and U. Pöschl. 2015. Ice nucleation activity in the widespread soil fungus Mortierella alpina. Biogeosciences 12:1057–1071.

Good, A. G., and P. H. Beatty. 2011. Fertilizing nature: a tragedy of excess in the commons. PLoS biology 9:e1001124.

Grządziel, J., and A. Gałązka. 2019. Fungal biodiversity of the most common types of polish soil in a long-term microplot experiment. Frontiers in Microbiology 10:6.

Guo, J., W. Liu, C. Zhu, G. Luo, Y. Kong, N. Ling, M. Wang, J. Dai, Q. Shen, and S. Guo. 2018. Bacterial rather than fungal community composition is associated with microbial activities and nutrient-use efficiencies in a paddy soil with short-term organic amendments. Plant and soil 424:335–349.

Guo, J. H., X. J. Liu, Y. Zhang, J. L. Shen, W. X. Han, W. F. Zhang, P. Christie, K. Goulding, P. M. Vitousek, and F. Zhang. 2010. Significant acidification in major Chinese croplands. Science 327:1008–1010.

Harris, J. 2009. Soil microbial communities and restoration ecology: facilitators or followers? Science 325:573–574.

Hartmann, M., B. Frey, J. Mayer, P. Mäder, and F. Widmer. 2015. Distinct soil microbial diversity under long-term organic and conventional farming. The ISME journal 9:1177–1194.

Hayat, R., S. Ali, U. Amara, R. Khalid, and I. Ahmed. 2010. Soil beneficial bacteria and their role in plant growth promotion: a review. Annals of microbiology 60:579–598.

Hu, X., J. Liu, D. Wei, P. Zhu, X. a. Cui, B. Zhou, X. Chen, J. Jin, X. Liu, and G. Wang. 2017. Effects of over 30-year of different fertilization regimes on fungal community compositions in the black soils of northeast China. Agriculture, Ecosystems & Environment 248:113–122.

Huang, F., Z. Liu, H. Mou, J. Li, P. Zhang, and Z. Jia. 2019. Impact of farmland mulching practices on the soil bacterial community structure in the semiarid area of the loess plateau in China. European Journal of Soil Biology 92:8–15.

Huang, R., Y. Wang, J. Liu, J. Gao, Y. Zhang, J. Ni, D. Xie, Z. Wang, and M. Gao. 2020. Partial substitution of chemical fertilizer by organic materials changed the abundance, diversity, and activity of nirS-type denitrifying bacterial communities in a vegetable soil. Applied Soil Ecology 152:103589.

Jannoura, R., C. Bruns, and R. G. Joergensen. 2013. Organic fertilizer effects on pea yield, nutrient uptake, microbial root colonization and soil microbial biomass indices in organic farming systems. European Journal of Agronomy 49:32–41.

Jiang, C., P. Liu, M. Liu, M. Wu, and Z. Li. 2017. Dynamics of aggregates composition and C, N distribution in rhizosphere of rice plants in red paddy soils different in soil fertility. Acta Pedologica Sinica 54:138–149.

Kumar, U., A. K. Nayak, M. Shahid, V. V. Gupta, P. Panneerselvam, S. Mohanty, M. Kaviraj, A. Kumar, D. Chatterjee, and B. Lal. 2018. Continuous application of inorganic and organic fertilizers over 47 years in paddy soil alters the bacterial community structure and its influence on rice production. Agriculture, Ecosystems & Environment 262:65–75.

Kumar, U., P. Panneerselvam, V. Govindasamy, L. Vithalkumar, M. Senthilkumar, A. Banik, and K. Annapurna. 2017. Long-term aromatic rice cultivation effect on frequency and diversity of diazotrophs in its rhizosphere. Ecological Engineering 101:227–236.

Kwak, M.-J., H. G. Kong, K. Choi, S.-K. Kwon, J. Y. Song, J. Lee, P. A. Lee, S. Y. Choi, M. Seo, and H. J. Lee. 2018. Rhizosphere microbiome structure alters to enable wilt resistance in tomato. Nature biotechnology 36:1100–1109.

Legrand, F., A. Picot, J. F. Cobo-Díaz, M. Carof, W. Chen, and G. Le Floch. 2018. Effect of tillage and static abiotic soil properties on microbial diversity. Applied Soil Ecology 132:135–145.

Li, B., L. Guo, H. Wang, Y. Li, H. Lai, X. Wang, and X. Wei. 2021. Bio-Organic Fertilizers Manipulate Abundance Patterns of Rhizosphere Soil Microbial Community Structure To Improve Tomato Productivity.

Liu, M., F. Hu, X. Chen, Q. Huang, J. Jiao, B. Zhang, and H. Li. 2009. Organic amendments with reduced chemical fertilizer promote soil microbial development and nutrient availability in a subtropical paddy field: the influence of quantity, type and application time of organic amendments. Applied Soil Ecology 42:166–175.

Liu, X.-Y., J.-D. Zou, L.-L. Xu, X.-Y. Zhang, F.-T. Yang, X.-Q. Dai, Z.-Q. Wang, and X.-M. Sun. 2014. Effects of different fertilizer species on carbon and nitrogen leaching in a reddish paddy soil. Huan jing ke xue= Huanjing kexue 35:3083–3090.

Lyu, J., L. Jin, N. Jin, J. Xie, X. Xiao, L. Hu, Z. Tang, Y. Wu, L. Niu, and J. Yu. 2020. Effects of different vegetable rotations on fungal community structure in continuous tomato cropping matrix in greenhouse. Frontiers in Microbiology 11:829.

Marschner, P., E. Kandeler, and B. Marschner. 2003. Structure and function of the soil microbial community in a long-term fertilizer experiment. Soil Biology and Biochemistry 35:453–461.

Martínez-García, L. B., G. Korthals, L. Brussaard, H. B. Jørgensen, and G. B. De Deyn. 2018. Organic management and cover crop species steer soil microbial community structure and functionality along with soil organic matter properties. Agriculture, Ecosystems & Environment 263:7–17.

Mendes, R., M. Kruijt, I. De Bruijn, E. Dekkers, M. van der Voort, J. H. Schneider, Y. M. Piceno, T. Z. DeSantis, G. L. Andersen, and P. A. Bakker. 2011. Deciphering the rhizosphere microbiome for disease-suppressive bacteria. Science 332:1097–1100.

Mukta, J. A., M. Rahman, A. A. Sabir, D. R. Gupta, M. Z. Surovy, M. Rahman, and M. T. Islam. 2017. Chitosan and plant probiotics application enhance growth and yield of strawberry. Biocatalysis and Agricultural Biotechnology 11:9–18.

Oksanen, J., F. G. Blanchet, M. Friendly, R. Kindt, P. Legendre, D. McGlinn, P. R. Minchin, R. O’Hara, G. Simpson, and P. Solymos. 2020. vegan: Community Ecology Package. R package version 2.5-6. 2019.

Philippot, L., J. M. Raaijmakers, P. Lemanceau, and W. H. Van Der Putten. 2013. Going back to the roots: the microbial ecology of the rhizosphere. Nature Reviews Microbiology 11:789–799.

Pieterse, C. M., R. de Jonge, and R. L. Berendsen. 2016. The soil-borne supremacy. Trends in plant science 21:171–173.

Raja, N. 2013. Biopesticides and biofertilizers: ecofriendly sources for sustainable agriculture. J Biofertil Biopestici 4:1–2.

Savci, S. 2012. An agricultural pollutant: chemical fertilizer. International Journal of Environmental Science and Development 3:73.

Semida, W. M., H. R. Beheiry, M. Sétamou, C. R. Simpson, T. A. Abd El-Mageed, M. M. Rady, and S. D. Nelson. 2019. Biochar implications for sustainable agriculture and environment: A review. South African Journal of Botany 127:333–347.

Sharma, S. K., A. Ramesh, M. P. Sharma, O. P. Joshi, B. Govaerts, K. L. Steenwerth, and D. L. Karlen. 2010. Microbial community structure and diversity as indicators for evaluating soil quality. Pages 317–358 Biodiversity, biofuels, agroforestry and conservation agriculture. Springer.

Shen, W., Y. Ni, N. Gao, B. Bian, S. Zheng, X. Lin, and H. Chu. 2016. Bacterial community composition is shaped by soil secondary salinization and acidification brought on by high nitrogen fertilization rates. Applied Soil Ecology 108:76–83.

Simonin, M., N. Nunan, J. M. Bloor, V. Pouteau, and A. Niboyet. 2017. Short-term responses and resistance of soil microbial community structure to elevated CO2 and N addition in grassland mesocosms. FEMS microbiology letters 364:fnx077.

Tamreihao, K., D. S. Ningthoujam, S. Nimaichand, E. S. Singh, P. Reena, S. H. Singh, and U. Nongthomba. 2016. Biocontrol and plant growth promoting activities of a Streptomyces corchorusii strain UCR3-16 and preparation of powder formulation for application as biofertilizer agents for rice plant. Microbiological research 192:260–270.

Team, R. C. 2018. R: A language and environment for statistical computing.

Trivedi, P., M. Delgado-Baquerizo, T. C. Jeffries, C. Trivedi, I. C. Anderson, K. Lai, M. McNee, K. Flower, B. Pal Singh, and D. Minkey. 2017. Soil aggregation and associated microbial communities modify the impact of agricultural management on carbon content. Environmental microbiology 19:3070–3086.

Van Der Heijden, M. G., R. D. Bardgett, and N. M. Van Straalen. 2008. The unseen majority: soil microbes as drivers of plant diversity and productivity in terrestrial ecosystems. Ecology letters 11:296–310.

Wang, P., H. Wang, Y. Qiu, L. Ren, and B. Jiang. 2018a. Microbial characteristics in anaerobic digestion process of food waste for methane production–A review. Bioresource technology 248:29–36.

Wang, Y., H. Ji, Y. Hu, R. Wang, J. Rui, and S. Guo. 2018b. Different selectivity in fungal communities between manure and mineral fertilizers: a study in an alkaline soil after 30 years fertilization. Frontiers in Microbiology 9:2613.

Wei, M., G. Hu, H. Wang, E. Bai, Y. Lou, A. Zhang, and Y. Zhuge. 2017. 35 years of manure and chemical fertilizer application alters soil microbial community composition in a Fluvo-aquic soil in Northern China. European Journal of Soil Biology 82:27–34.

Xiong, W., S. Guo, A. Jousset, Q. Zhao, H. Wu, R. Li, G. A. Kowalchuk, and Q. Shen. 2017. Bio-fertilizer application induces soil suppressiveness against Fusarium wilt disease by reshaping the soil microbiome. Soil Biology and Biochemistry 114:238–247.

Xiong, W., A. Jousset, S. Guo, I. Karlsson, Q. Zhao, H. Wu, G. A. Kowalchuk, Q. Shen, R. Li, and S. Geisen. 2018. Soil protist communities form a dynamic hub in the soil microbiome. The ISME journal 12:634–638.

Yan, M., S. Chen, T. Huang, B. Li, N. Li, K. Liu, R. Zong, Y. Miao, and X. Huang. 2020. Community compositions of phytoplankton and eukaryotes during the mixing periods of a drinking water reservoir: dynamics and interactions. International journal of environmental research and public health 17:1128.

Yang, X., L. Chen, X. Yong, and Q. Shen. 2011. Formulations can affect rhizosphere colonization and biocontrol efficiency of Trichoderma harzianum SQR-T037 against Fusarium wilt of cucumbers. Biology and fertility of soils 47:239–248.

Yao, Q., J. Liu, Z. Yu, Y. Li, J. Jin, X. Liu, and G. Wang. 2017. Three years of biochar amendment alters soil physiochemical properties and fungal community composition in a black soil of northeast China. Soil Biology and Biochemistry 110:56–67.

Ye, L., X. Zhao, E. Bao, J. Li, Z. Zou, and K. Cao. 2020. Bio-organic fertilizer with reduced rates of chemical fertilization improves soil fertility and enhances tomato yield and quality. Scientific reports 10:1–11.

Yelle, D. J., J. Ralph, F. Lu, and K. E. Hammel. 2008. Evidence for cleavage of lignin by a brown rot basidiomycete. Environmental microbiology 10:1844–1849.

Zhang, Q.-C., I. H. Shamsi, D.-T. Xu, G.-H. Wang, X.-Y. Lin, G. Jilani, N. Hussain, and A. N. Chaudhry. 2012. Chemical fertilizer and organic manure inputs in soil exhibit a vice versa pattern of microbial community structure. Applied Soil Ecology 57:1–8.

Zhang, X., W. Liu, G. Zhang, L. Jiang, and X. Han. 2015. Mechanisms of soil acidification reducing bacterial diversity. Soil Biology and Biochemistry 81:275–281.

Zhang, Y., X. Hao, T. W. Alexander, B. W. Thomas, X. Shi, and N. Z. Lupwayi. 2018. Long-term and legacy effects of manure application on soil microbial community composition. Biology and fertility of soils 54:269–283.

Zhang, Y., C. Li, Y. Wang, Y. Hu, P. Christie, J. Zhang, and X. Li. 2016. Maize yield and soil fertility with combined use of compost and inorganic fertilizers on a calcareous soil on the North China Plain. Soil and Tillage Research 155:85–94.

Zhang, Y., H. Shen, X. He, B. W. Thomas, N. Z. Lupwayi, X. Hao, M. C. Thomas, and X. Shi. 2017. Fertilization shapes bacterial community structure by alteration of soil pH. Frontiers in Microbiology 8:1325.

Zhao xiang, W., L. Huihu, L. Qiaoli, Y. Changyan, and Y. Faxin. 2020. Application of bio-organic fertilizer, not biochar, in degraded red soil improves soil nutrients and plant growth. Rhizosphere 16:100264.

Zheng, X., Y. Zhu, Z. Wang, H. Zhang, M. Chen, Y. Chen, J. Wang, and B. Liu. 2020. Effects of a novel bio-organic fertilizer on the composition of rhizobacterial communities and bacterial wilt outbreak in a continuously mono-cropped tomato field. Applied Soil Ecology 156:103717.

Zhu, Y.-G., J.-Q. Su, Z. Cao, K. Xue, J. Quensen, G.-X. Guo, Y.-F. Yang, J. Zhou, H.-Y. Chu, and J. M. Tiedje. 2016. A buried Neolithic paddy soil reveals loss of microbial functional diversity after modern rice cultivation. Science Bulletin 61:1052–1060.

